# Generation of a Transgenic Mouse Model for Investigating Mitochondria in Sperm

**DOI:** 10.1101/2024.12.08.627442

**Authors:** Hironmoy Sarkar, Suryaprakash R Batta, Neerja Wadhwa, Subeer S Majumdar, Bhola Shankar Pradhan

**Author notes:** Correspondence: author: Hironmoy Sarkar, Bhola Shankar Pradhan.

## Abstract

Mitochondria play a crucial role in the sperm development; however, the mechanisms regulating their function in sperm remain poorly understood. Developing a method to regulate the expression of a target gene within the mitochondria of sperm is a vital step in this area of research. In this study, we aimed to create a system for expressing a transgene in the mitochondria of sperm. As a proof of concept, we generated transgenic mice that express green fluorescent protein (GFP) fused with a mitochondrial localization signal (MLS) driven by the phosphoglycerate kinase 2 (PGK2) promoter, which facilitates the transgene expression in the sperm. Although the PGK2 promoter has previously shown to drive gene expression in spermatocytes and spermatids, the novelty of our approach lies in the combination of PGK2-driven MLS-GFP expression to study mitochondria *in vivo*. We established two founder lines of transgenic mice through pronuclear microinjection, and MLS-GFP expression was confirmed in the mitochondria of sperm cells using fluorescence microscopy and flow cytometry. Consequently, we provide a novel platform for investigating mitochondrial function in sperm, where GFP can be substituted with other genes of interest to examine their effects on mitochondria. This system specifically targets sperm mitochondria, offering an innovative approach for studying mitochondrial function *in vivo*.

## Introduction

The rise in the cases of infertility has become a global concern recently, and around 7% of the male population worldwide are infertile (Matzuk and Lamb, 2008; Sharpe, 2011; Hentrich et al., 2011; Thirumavalavan et al., 2019). The dysfunction of mitochondria contributes significantly towards the male infertility (Tesarik et al., 2023). Mitochondria are dynamic organelles and play a crucial role in the metabolism of spermatozoa. In the spermatozoa, approximately 50-80 mitochondria are present in the midpiece (Freitas et al., 2017). They are implicated in various cellular functions such as oxidative phosphorylation and calcium homeostasis which are important for function of sperm (Aitken et al., 2012; Amaral et al., 2013; Boguenet et al., 2021). The mitochondria in spermatozoa regulate important functions relating to human fertility (Spinelli et al., 2018; Fu et al., 2021). The ROS-mediated damage to the spermatozoa contributes to 30% to 80% (approximately) of male infertility (Agarwal et al., 2014) and around 60% of idiopathic male infertility is associated with reduced sperm motility (Liu et al., 2015). The genetic modification of mitochondria and its role in sperm development has gained significant attention in recent years. The understanding of the function of mitochondria in sperm cells may provide critical insights into the molecular basis of male infertility (Park et al., 2021; Freitas et al., 2017).

Recent reports suggested that male infertility is associated with a loss of mitochondrial proteins in spermatozoa, which induces low sperm motility, reduces OXPHOS activity, and results in male infertility (Park et al., 2021). The reduction in the motility of the spermatozoa in cases of asthenozoospermia is associated with reduced mitochondrial respiratory activities and reduced mitochondrial membrane potential (Ruiz-Pesini et al., 1998; Piasecka et al., 2003). The inhibition of the protein synthesis machinery of the mitochondria of the spermatozoa leads to a reduction in sperm motility (Gur et al., 2006). Moreover, the interaction of mitochondrial fusion protein 2 (MFN2) and mitostatin-related protein 1 (MNS1), which regulate the biogenesis of sperm tail contributes towards sperm motility (Vadnais et al., 2014).

Despite the advancements in understanding the role of mitochondrial dynamics in the function of the cells (Otera et al., 2013; Wakai et al., 2014; Sandoval et al., 2014; Bader et al., 2020), the molecular mechanisms regulating this process during spermatogenesis, particularly in the development of spermatozoa, are not fully understood. Importantly, the techniques to study the roles of various genes affecting the mitochondria of the spermatozoa *in vivo* are limited. Therefore, there is an urgent need to develop a method to target the gene specifically to the mitochondria of the spermatozoa *in vivo*.

The dynamics of mitochondria in spermatozoa, including their distribution, fusion, and fission, are important for sperm function and viability. During the development of sperm, mitochondria undergo significant structural and functional changes. The role of mitochondrial fusion and fission in sperm motility is increasingly understood, as proper mitochondrial dynamics are necessary for the motility and structural integrity of the sperm tail (Green et al., 2022; Raad M et al., 2024). The use of various staining dyes to label the mitochondria of the spermatozoa, such as Mitotracker, leads to the defect in the fertilizing ability of spermatozoa *in vitro*, highlighting the need for alternative approaches. Therefore, fluorescent proteins may be used for studying the effect of mitochondria in the development of the spermatozoa *in vivo*.

Previously, the PGK2 promoter has been demonstrated as a testis-specific promoter (Zhang et al., 1999). The PGK2 promoter drives expression of transgenes in spermatocytes and spermatids *in vivo* (Kramer et al., 1981; Danshina et al., 2010; McCarrey et al., 1992; Yoshioka et al., 2007). In this study, we seek to develop a novel method for studying the mitochondria within spermatozoa development by combining the GFP with a mitochondrial localization signal (MLS) along with PGK2 promoter. We hypothesize that the PGK2 promoter can drive the expression of GFP-tagged MLS in the mitochondria of the spermatozoa *in vivo*.

This approach not only enhances our understanding of mitochondrial behavior during spermatogenesis but also establishes a foundation for exploring the molecular basis of male infertility related to mitochondrial dysfunction. Although the role of PGK2 promoter in spermatogenesis is well established, its use for targeting the gene expression in the mitochondria of the spermatozoa is not demonstrated so far. In this study, we used the PGK2 promoter driving the expression of GFP tagged to a mitochondrial localization signal (PGK2-MLS-GFP) to express GFP specifically in the spermatozoa of transgenic mice.

## Material and methods

### Animals

All the mice used for the present study were of the FVB/J strain. The mice were obtained from the small animal facility of the National Institute of Immunology. The mice were housed in a climate-controlled facility under standard light (14-h light-10-h dark cycle), temperature (23 °C), and humidity (50±5%). The mice were housed in individually ventilated cages with (IVC) ad libitum access to acidified autoclaved water and handled by trained personnel. Mice were used as per the guidelines laid down by the CPCSEA (Committee for the Purpose of Control and Supervision of the Experiments on Animals). Protocols for experiments were approved by the Institutional Animal Ethics Committee (IAEC) of the National Institute of Immunology, India, constituted by CPCSEA (IAEC number: IAEC 249/10). Mice were euthanized by cervical dislocation as approved by the Institutional Animal Ethics Committee of the National Institute of Immunology.

### Transgene constructs

A total of three constructs (EGFPN2, CMV_MLS-EGFP and PGK2_MLS-EGFP) were used in this study. The schematic diagrams of the constructs were provided in **Fig 1A**. The constructs for mitochondrial gene targeting were generated through standard molecular cloning methods as described previously (Pradhan et al., 2016). Plasmid DNA was isolated from overnight-grown culture of E. coli (dh5α) using a plasmid DNA isolation kit (Advanced Micro Devices, India) following the manufacture’s recommendation. For the generation of CMV_MLS-EGFP construct, the mitochondrial localization signal of the *COXVIII* gene of murine origin (*TCCGTCCTGACGCCGCTGCTGCTGCGGGGCTT-GACAGGCTCGGCCCGGCGGCTCCCAGTGCCGCGCGCCAAGA*) was amplified from mouse genomic DNA using primers MLSF: *GCCGCCAAGCTTGTGTACTCCGTGCCATCATG* and MLSR: *GGCGTGGATCCCCGGCGGCAACGAATGGA*. The PCR was performed using pfu DNA polymerase (Fermentas, Pittsburgh, PA, USA), following standardized thermal cycling parameters (initial denaturation at 95 °C for 2 min, denaturation at 95 °C for 30 s, annealing at 60 °C for 30 s, and extension at 68 °C for 30 s for 35 cycles, followed by final extension for 10 min at 72 °C). The amplicon (135 bp) was cloned between the two restriction enzyme sites HindIII and BamHI of the EGFPN2 vector. For the generation of PGK2_MLS-EGFP construct, the PGK2 promoter was amplified from mouse genomic DNA using primers PGKF: *GTCGT ATTAATCTAGAAATAGGATTGATGAC* and PGKR: *GTCGTCCTCGAGCTTGACAATATAAAGACATA*. The PCR was performed using pfu DNA polymerase (Fermentas, Pittsburgh, PA, USA), following standardized thermal cycling parameters (initial denaturation at 95 °C for 2 min, denaturation at 95 °C for 30 s, annealing at 60 °C for 30 s, and extension at 68 °C for 2 min for 35 cycles, followed by final extension for 10 min at 72 °C). The amplicon (1467 bp) was cloned between the two restriction enzyme sites, AseI and XhoI, of the CMV_MLS-EGFP vector by substituting the CMV promoter with the PGK2 promoter. The sequences of the construct are provided in the **Supplementary Fig. 3**.

**Figure 1.**
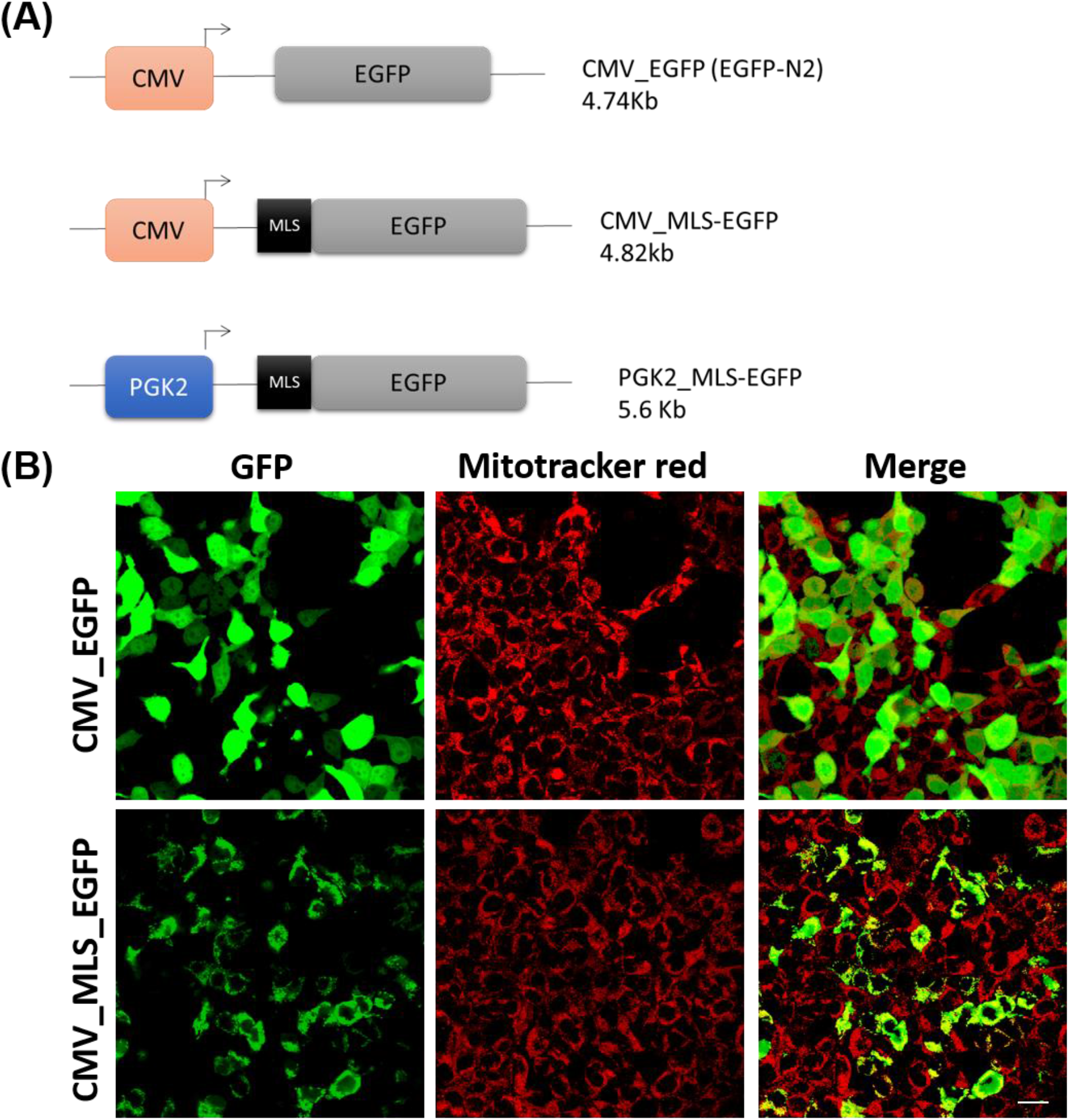
Validation of MLS sequence in HEK293 cells. (A) A schematic diagram of the constructs used in this study. (B) HEK 293 cells were transfected with CMV_EGFP (control) and CMV_MLS-EGFP; and co-stained with Mitotracker red dye. The images were taken 24h after transfection. The left panel was expression for GFP, middle panel was for the mitotracker of the same cells, and the right panel was the co-localization of GFP and Mitotracker red. Scale bar: 10µm.

### Cell culture and transfection

The human embryonic kidney (HEK293) cells (CRL-1573, ATCC) were a kind gift from Dr. A.C. Banerjea, National Institute of Immunology, New Delhi. The HEK293 cells were cultured in Dulbecco’s Modified Eagle’s Medium (DMEM) (670087, Gibco™) containing 10% fetal bovine serum supplemented with penicillin, streptomycin, and Fungizone at 37°C in the presence of 5% CO_2_. Transfection of HEK293 cells was performed as described by us previously (Pradhan et al., 2016). Transient transfections with CMV_MLS-EGFP plasmid were performed with Lipofectamine 2000 (11668019, Invitrogen), according to the manufacturer’s instructions. For the control, we used EGFPN2 (CMV_EGFP) plasmid, which lacks the mitochondrial localization signal. After 24 h, the transfected cells were stained with 20 nM of the mitochondrial-specific dye, MitoTracker Red CMXRos (M7512, Thermo Fisher Scientific), for 15 min at 37 °C, followed by three washes with 1X PBS.

### Pronuclear microinjection of transgene cassettes and oviductal embryo transfer

This method was performed as described previously (Sarkar et al., 2020). Transgenic mice were generated via pronuclear microinjection (Auerbach et al., 2004; Brinster et al., 1981; Costantini et al., 1981; Gordon et al., 1980). For pronuclear microinjection, female mice were superovulated using PMSG and hCG followed by cohabitation with male mice. Oviducts were collected from donor mice (euthanized by cervical dislocation) and placed into Brinster’s Modified Oocyte Culture Media (BMOC) containing hyaluronidase (1 mg/mL). Embryos were collected and transferred to the 60 mm dish containing 100 µL drops of BMOC overlaid with mineral oil. The linearized transgene cassette (PGK2-MLS-EGFP) was prepared at a final concentration of 4 ng/µL for microinjection. DNA was microinjected into the male pronucleus of fertilized eggs using a Narishighe micromanipulator. Each manipulated embryo was then transferred into the pre-incubated BMOC-containing dish and maintained at 37°C in a 5% CO_2_ incubator. Embryos were incubated till the two-cell stage. Two pseudo-pregnant female mice were generated by mating with vasectomized male mice. 2-cell stage embryos were transferred into the ampulla of the oviduct of these pseudo-pregnant recipient mice. Approximately 20 microinjected embryos were transferred in the oviductal ampulla. After the gestation period (21 ± 2 days), recipient pups were born.

### Establishment of Transgenic Lines

We obtained many pups from the two pseudo-pregnant recipient mice after the embryo transfer. The pups born were screened through PCR as described by us previously (Pradhan et al., 2016; Usmani et al., 2013). For this, the genomic DNA was isolated from the tail biopsy of these mice (around 21 days of age) by proteinase K lysis and organic extraction with phenol-chloroform. The genotyping was performed by PCR using Taq polymerase and the following primers: MKSF: *GTGTACTCCGTGCCATCATG* and GFPR: *TGCCGTAGGTGGCATCGCCC*; following standardized thermal cycling parameters (initial denaturation at 95 °C for 2 min, denaturation at 95 °C for 30 s, annealing at 60 °C for 30 s and extension at 72 °C for 30 s for 35 cycles, followed by final extension for 10 min at 72 °C). The PCR products were analyzed by TAE agarose gel electrophoresis. The amplicon size of 258 bp indicates the transgene positive mice. As a loading control, PCR for mice gene *peptidylprolyl isomerase A (Ppia*) was amplified from each tail biopsy using the primer PPIAF: *GTCGCTTTTCGCCGCTTGC* and PPIAR: *TCTGCTGTCTTTGGAACT* fol-lowing standardized thermal cycling parameters (initial denaturation at 95 °C for 2 min, denaturation at 95 °C for 30 s, annealing at 60 °C for 30 s, and extension at 72 °C for 30 s for 35 cycles, followed by final extension for 10 min at 72 °C). The amplicon size (120 bp) suggests good quality of genomic DNA. The founder mice were mated with either wildtype females or wild-type males to establish the separate transgenic lines, and the transgene integration in the progeny was evaluated by PCR. The litter size of the control and PGK2-MLS-GFP mice was analyzed.

### Sperm Collection

The sperm cells were collected from four PGK2-MLS-GFP transgenic male mice and four wild-type male mice as described previously (Basu et al., 2018; Sarkar et al., 2021). These mice were euthanized at around 90 days of age. The entire intact epididymis was surgically removed from these mice and transferred into a 1.5 ml microcentrifuge tube containing 1 ml of 1X PBS and ruptured thereafter for sperm cell collection. The sperm cells of the control and PGK2-MLS-GFP mice were analyzed for motility under a light microscope. For this, around 100 sperm cells were counted per field, and ten different fields were counted for each sample. A fraction of these samples was analyzed in fluorescence microscopy, and another fraction was analyzed in a flow cytometer for the detection of GFP.

### Fluorescence Microscopy

The HEK293 cells transfected with CMV_EGFP plasmid (control) or with CMV_MLS-EGFP plasmid were analyzed under fluorescent microscope for GFP expression. The mitochondria of these cells were visualized by staining with MitoTracker Red CMXRos. Images were acquired from five different fields, and experiments were performed in triplicates.

The sperm cells from the PGK2-MLS-GFP transgenic male mice (n = 4) and wild-type male mice (n = 4) were analyzed for the presence of GFP under bright field and ultraviolet illumination (with FITC and BLUE/CYAN filter) using a Nikon Eclipse TE2000-S inverted microscope (Nikon Corporation, Chiyoda-ku, Tokyo, Japan) attached to a DS-5 M camera assisted by Digital Sight DS-L1 software for capturing the images. Around 100 sperm cells were analyzed for each mouse for the presence of GFP.

### Flow Cytometry Analysis

The sperm cells from the PGK2-MLS-GFP transgenic male mice (n = 4) and wild-type male mice (n = 4) were analyzed for the presence of GFP in a flow cytometer (FACS caliber, BD Biosciences). Around 10,000 sperm cells from each mouse were analyzed for the presence of GFP. Data were collected and analyzed using FlowJo software (TreeStar, Inc.) to assess the percentage of GFP-positive sperm.

### Statistical Analysis

All statistical analyses were performed using Graphpad Prism 5 (Graphpad Software, Lajolla, CA, USA). Data are represented as means ± SEM. Significance was determined by the Mann–Whitney test and a P value.

## Results

### Validation of MLS sequence in vitro

The effectiveness of the MLS sequence of the *COXVIII* gene of human in targeting GFP to mitochondria was demonstrated in HeLa cells before (Rizzuto et al.,1995; Rizzuto et al., 1996). Since we plan to target the gene expression specifically in the mitochondria of the spermatozoa of the mice *in vivo*, we used the MLS sequence of the *COXVIII* gene of mice. However, the sperm cells are difficult to transfect *in vitro*. So we tested the MLS sequence of *COXVIII* gene of mice in HEK293 cells, as these cells are comparatively easier to transfect. For this, we generated the CMV_MLS-EGFP construct, which contains the MLS sequence of the *COXVIII* gene of mice along with the CMV promoter, which is functional in HEK293 cells. To determine the effectiveness of MLS sequence in targeting the gene expression in mitochondria *in vitro*, we transfected the HEK293 cells with CMV_MLS-EGFP construct. As a control, we transfected the HEK293 cells with EGPN2 vector (which lacks the MLS sequence). We observed the tubular pattern of GFP signal in the HEK293 cells transfected with CMV_MLS-EGFP construct. Importantly, the signal for GFP in these cells was colocalized with the red signal of the mitochondria which were stained with MitoTracker Red CMXRos (mitochondria-specific dye) suggesting that MLS was functional (**Fig. 1B**). The control cells transfected with the EGFPN2 construct showed a cytoplasmic GFP signal that did not colocalize with the MitoTracker Red CMXRos. These results also suggest that the MLS sequence of the *COXVIII* gene of mice is functional in HEK293 cells.

### Generation of Transgenic Mice

Since the MLS sequence of the *COXVIII* gene of mice is effective in targeting gene expression in the mitochondria in vitro, we used this MLS sequence for testing its functionality in the spermatozoa of mice. Previously, the tissue-specific activity of the PGK2 promoter is demonstrated to drive the gene expression of the post-meiotic genes in the male germline, particularly in the spermatozoa (Zhang et al., 1999). So we generated a PGK2_MLS-GFP construct by replacing the CMV promoter with the PGK2 promoter of mice in the CMV_MLS-EGFP construct. Using the classical pronuclear microinjection method, we inserted the PGK2_MLS-GFP construct into the pronucleus of the embryos and transferred these embryos into the pseudo-pregnant females. A schematic diagram of the generation of the transgenic line is presented in **Fig. 2**. We obtained many pups from the two recipient females after the pronuclear microinjection. The pups born were screened for the integration of transgenes using the genomic DNA of tail biopsies by PCR. We observed that many mice were positive for integration of transgene (PCR product size: 258 bp) (**Fig. 3**). As a loading control, we used the PCR for *Ppia*. Further, we selected two founder mice out of these for establishing the transgenic line expressing PGK2-MLS-GFP by mating them with the wild-type males or female mice. The pups born from this mating were screened for PCR, and the transgenic mice were used for further study. The litter size from the PGK2-MLS-GFP mice was compared with that of wild-type mice (control). We did not observe any significant (p ≤ 0.05) difference in the litter size of PGK2-MLS-GFP mice as compared to that of the control suggesting that PGK2-MLS-GFP mice were fertile (**Suppl Fig. 2**).

**Figure 2.**
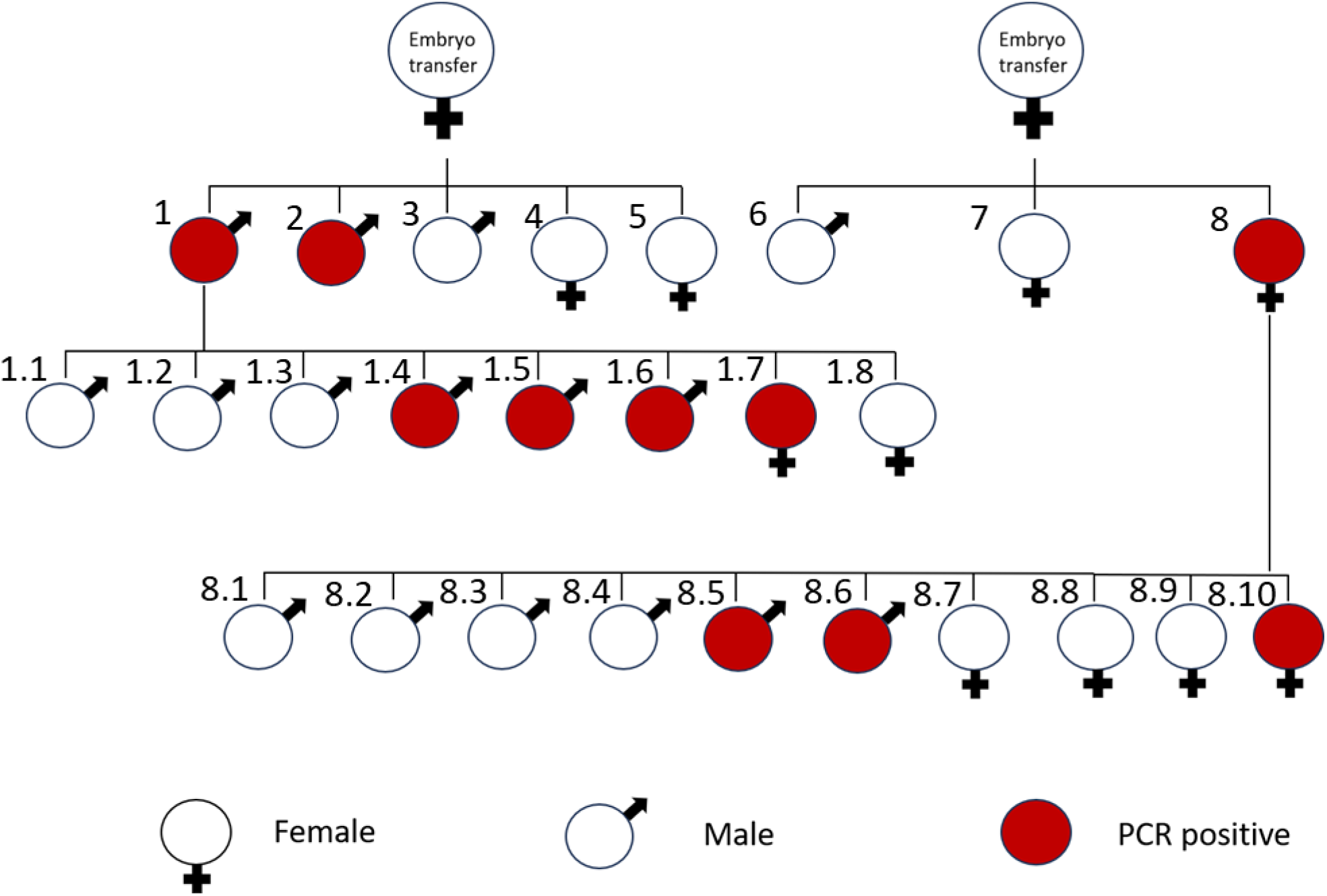
A schematic diagram of generation of PGK2-MLS-GFP transgenic mice after embryo transfer. The male and female mice were indicated in the figure. The PCR positive progenies were marked in red. After the embryo transfer, mice No. 1,2, and 8 were transgene positive (founder). The founder mice No. 1 and No. 8 with wild type mice, the mice No 1.4, 1.5, 1.6, 1.7, 8.5, 8.6, & 8.10 were detected to be PCR positive for transgene integration.

**Figure 3.**
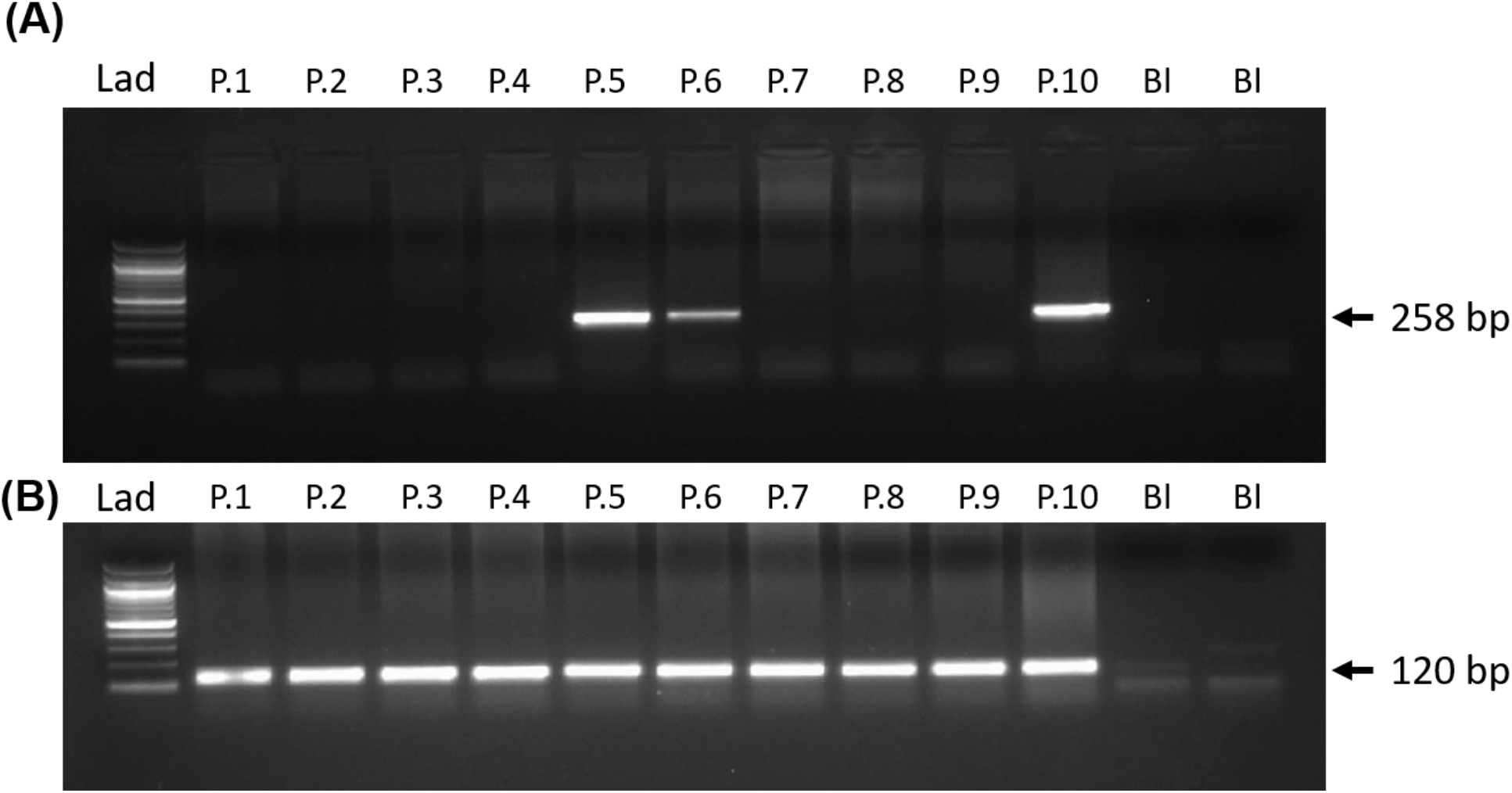
Generation of PGK2-MLS-GFP transgenic mice. Detection of transgene by PCR. PCR screening of progeny for generation of PGK2-MLS-GFP mice. (A) A band size of 258 bp represented the transgene positive mice. (B) PCR of the gene *Ppia* (120 bp) was used to determine the quality of the isolated gDNA from mice (loading control). Lad: 100 bp DNA ladder; Bl = blank containing only PCR master mix, P.1 to P.10 indicated the progeny numbers in the gel.

### Localization of the GFP in the mitochondria of spermatozoa in vivo

Since the PGK2 promoter drives expression of transgenes in spermatocytes and spermatids *in vivo* (Kramer et al., 1981; Danshina et al., 2010; McCarrey et al., 1992; Yoshioka et al., 2007), we evaluated the sperm of the PGK2-MLS-GFP transgenic mice. We collected the sperm of the transgenic mice (n=4) and control mice (n=4). These live sperm cells (n=100) were observed under the fluorescence microscope for the presence of GFP. We observed the GFP signal in the spermatozoa collected from transgenic mice but not from the control mice (**Fig. 4**). The mitochondria are localized in the midpiece of the spermatozoa. Since we used the MLS sequence of the *COXVIII* gene of mice, we evaluated the mid-piece region of the GFP-positive spermatozoa under a fluorescence microscope. We observed the GFP signal in the midpiece of the spermatozoa of the transgenic mice (**Fig. 4 and Suppl Fig. 1**). There was no signal for GFP either in the head or tail region of the spermatozoa, suggesting that GFP was expressed specifically in the midpiece (where mitochondria are localized) of the spermatozoa of PGK2-MLS-GFP transgenic mice. Moreover, the motility of sperm was analyzed and there was no significant (p≤0.05) difference between the control and transgenic mice in terms of sperm motility suggesting that GFP expression did not affect the function of the sperm cells (**Suppl Fig. 2**).

**Figure 4.**
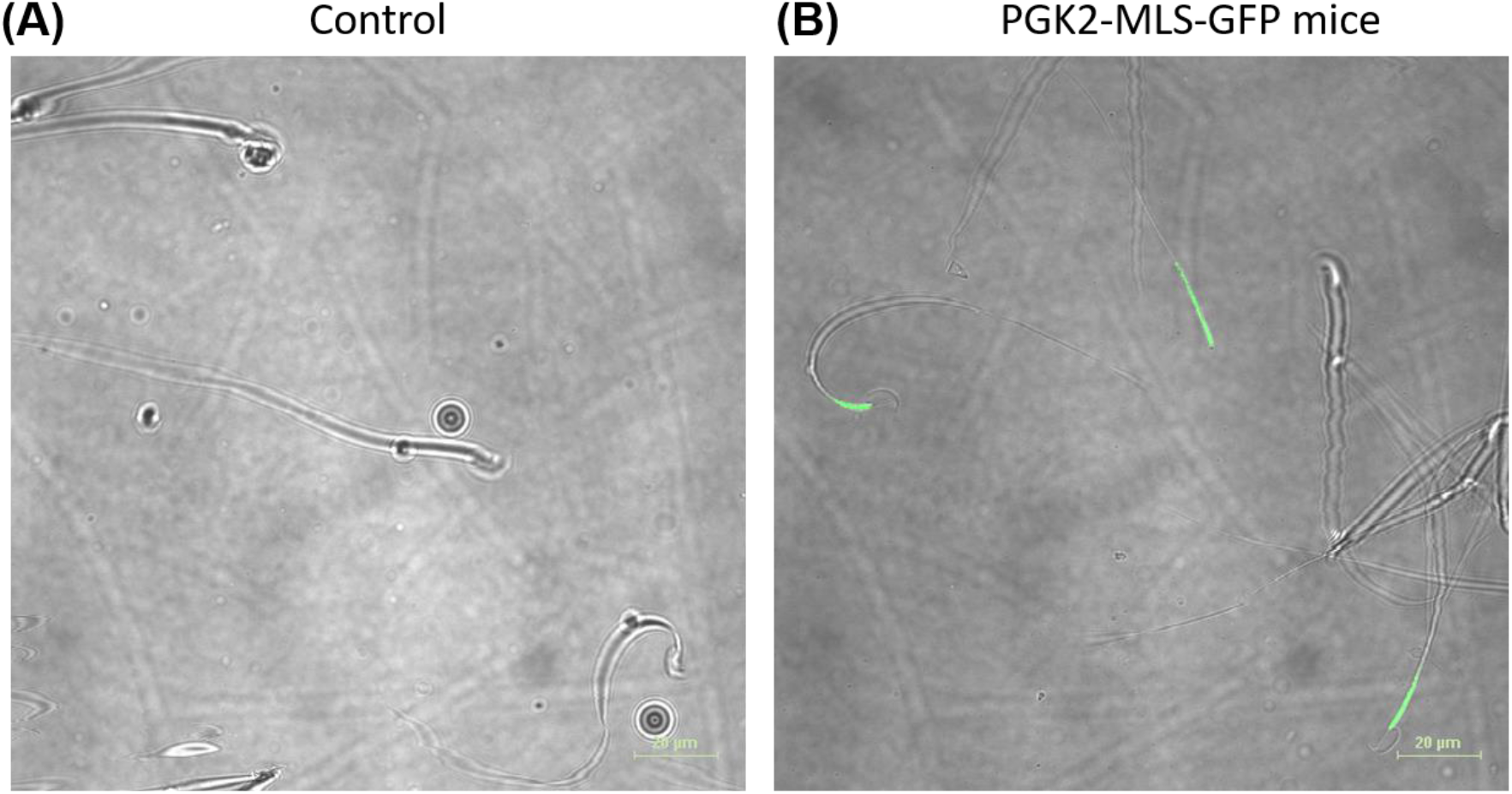
Detection of GFP-positive spermatozoa isolated from the epididymis of the PGK2-MLS-GFP transgenic mice by microscopy. Microscopic images of the sperm cells of wild-type mice (A) and transgenic mice (B). The GFP positive spermatozoa was detected by microscopy in the live sperm cells of the transgenic mice . Scale bar: 20µm.

### Analysis of GFP expression in the spermatozoa by Flow Cytometry (FACS)

Since we observed the GFP in the mitochondria of the spermatozoa of the PGK2-MLS-GFP transgenic mice by fluorescence microscopy, we performed the FACS analysis of sperm samples of these mice (n=4) to validate the microscopy data. As a control, we used the sperm cells of the wild-type mice (n=4). Around 10,000 cells per sample were analyzed for detection of GFP by flow cytometry. We observed a distinct shift in the sperm samples of PGK2-MLS-GFP transgenic mice, which suggests for the population of GFP-positive sperm (**Fig 5**). On the other hand, the sperm cells of the control wild-type mice did not show this shift, suggesting the absence of GFP-positive sperm. These results validate our observation from microscopic data *in vivo*. Moreover, these results along with the microscopy data, also suggest that the GFP expression was specifically in the mitochondria of the spermatozoa of the PGK2-MLS-GFP transgenic mice and the method was reproducible.

**Figure 5.**
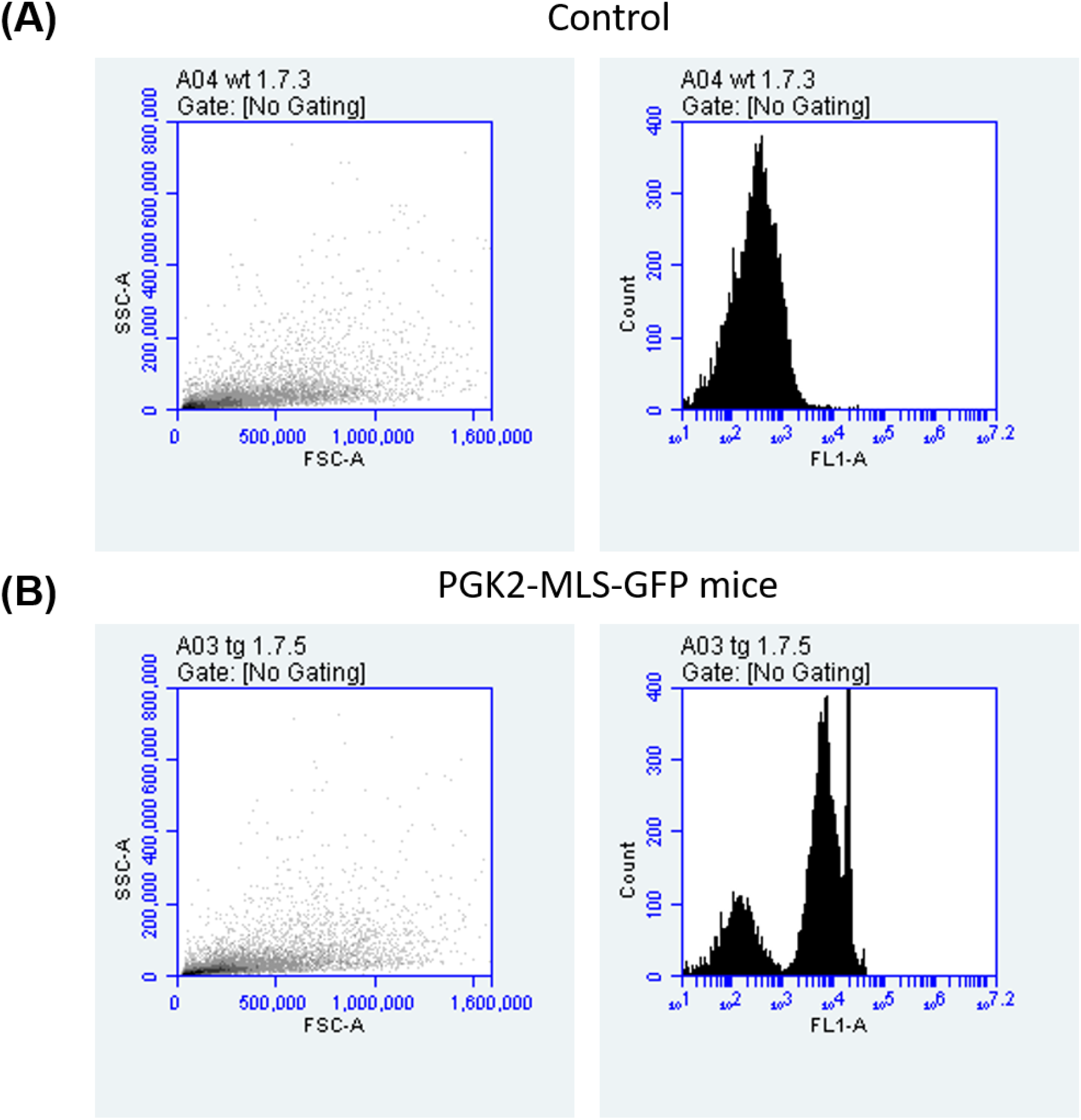
Detection of GFP-positive sperm cells collected from the epididymis of the PGK2-MLS-GFP transgenic mice by flow cytometry. (A) Scatter plot of one of the representative sperm samples and the corresponding histogram of wild-type mouse No 1.7.3. (B) Scatter plot of one of the representative sperm samples and the corresponding histogram of transgenic mouse No 1.7.5.

## Discussion

The present study was aimed to target the expression of transgene to the mitochondria of the spermatozoa *in vivo*. Here we demonstrated the targeting of GFP to the mitochondria of the spermatozoa of the transgenic mice as a proof of concept. For this, we used the construct containing the MLS sequence of the *COXVIII* gene and the PGK2 promoter of mice. This combination allows us to target the GFP specifically to the mitochondria of the spermatozoa. This conclusion is based on the following experimental data. First, we validated the effectiveness of the MLS sequence of the *COXVIII* gene in targeting GFP to the mitochondria in HEK293 cells. Secondly, we generated the PGK2-MLS-GFP transgenic mice and demonstrated the presence of GFP in the mitochondria of the spermatozoa by fluorescence microscopy and flow cytometry analysis. Our data also suggest the targeting of GFP in the midpiece of spermatozoa mitochondria did not affect its motility nor the fertility of the PGK2-MLS-GFP transgenic mice.

Previously, the MLS sequence of the *COXVIII* gene of both humans and mice was used for mitochondrial gene expression in HeLa cells and transgenic mice, respectively (Rizzuto et al., 1995; Rizzuto et al., 1996; Shitara et al., 2001; Wakai et al., 2014). In this study, we used the MLS of the *COXVIII* gene of mice and tested its effectiveness for mitochondrial gene targeting both in HEK293 cells and in transgenic mice, which suggests the reproducibility of the MLS sequence the *COXVIII* gene in mitochondrial gene targeting. Moreover, the MLS sequence of the *COXVIII* gene of mice is functional in human cells, such as HEK293 cells which suggests for the functionality of this sequence across species. Previous studies have utilized the constitutive promoter for GFP expression in the spermatozoa, which also led to expression of transgenes in other testicular cells such as Sertoli cells, Leydig cells etc. (Shitara et al., 2001; Barrasso et al., 2018; Yamaguchi et al., 2012). This strategy will not help in studying the function of a particular gene specifically in the spermatozoa. The specificity of the PGK2 promoter ensures minimal ectopic expression, which is critical for accurate functional analyses (Zhang et al., 1999). Thus, the use of PGK2 promoter, which is active only in meiotic spermatocytes and post-meiotic spermatids (Zhang et al., 1999) will address this limitation. In this study, we observed the GFP expression specifically in the spermatozoa, which is consistent with the previous studies. Further, we extended these findings by the integration of the MLS sequence along with PGK2 promoter, which enables precise targeting of GFP to the mitochondria of spermatozoa *in vivo*. This approach addresses a critical gap in male reproductive biology, providing insights into studying the role of mitochondria in spermatozoa development, motility, and fertilization.

Increasing evidence suggests the role of mitochondria in male infertility (Matilainen et al., 2017; Tesarik et al., 2023). Mitochondria in the midpieces of the spermatozoa are critical for energy production and motility and previous studies suggest a link between mitochondrial dysfunction and male infertility (Park et al., 2021). This approach can be used to study the effect of various mutations caused by environmental stress on mitochondrial functions, mitochondrial dynamics using advanced imaging techniques during spermatogenesis and the role of mitochondria in assisted reproduction technologies.

In this study, we demonstrated the functional importance of mitochondrial localization sequences in directing transgenes to the mitochondria of the sperm using a germline-specific promoter. Importantly, we confirmed that GFP expression in the mitochondria of spermatozoa did not affect its functions, as we did not find any difference in the sperm motility or reduction in offspring number from the male transgenic mice. This is in agreement with the previous study, which suggest that the expression of GFP in the mitochondria of the spermatozoa does not affect its function (Wang *et al*., *2023*).

This study may help in studies of mitochondrial health and dynamics in large populations of sperm cells (Park et al., 2021). Our study provides a platform to investigate mitochondrial biology during spermatogenesis. Mitochondria are vital for sperm function and is a major contributor to male infertility (Tesarik et al., 2023). Our studies will help to study the role of mitochondria in the development of the spermatozoa.

While our study demonstrates the feasibility of targeting transgenes to the mitochondria of the spermatozoa, we did not directly assess the mitochondrial biogenesis or role of various environmental stresses on mitochondrial function. Future studies will be needed to explore the role of various genes on the function of the mitochondria of the spermatozoa and thereby their role in affecting male fertility.

In conclusion, we establish a novel *in vivo* model for studying the role of transgenes in the mitochondria of the spermatozoa using a germline-specific promoter such as PGK2 and mitochondrial localization sequence of the *COXVIII* gene. These findings may have broad implications for mitochondrial research, particularly in male infertility and sperm development.

## Acknowledgments

We are thankful to all the staff of the Small Animal Facility. Thanks are due to Ram Singh, Dharamvir Singh and Birendar Roy for the technical assistance. We are grateful to the Director, NII for valuable support. We are grateful to Department of Biotechnology, Govt. of India, for providing the financial assistance under grants BT/PR11313/AAQ/01/376/2008, BT/HRD/35/01/01/2010 (TATA Innovation Award) and BT/PR10805/AAQ/1/576/2013. BSP was supported the Polish National Science Centre grant (2020/39/D/NZ5/02004). H.S. was supported by the Department of Biotechnology, Govt. of India grant (BT/PR46488/AAQ/1/886/2022). The authors declare that they have no competing interests.

## Author Contributions

Experiments were conceived and designed by B.S.P., H.S., and S.S.M. Experiments were performed by H.S., S.B., N.W. and B.SP. Data of the manuscript was analyzed by B.S.P, H.S. Manuscript was written and reviewed by B.S.P. and H.S. All authors have read and agreed to the published version of the manuscript.

## Institutional Review Board Statement

Mice were housed and used as per the guidelines laid down by the CPCSEA (Committee for the Purpose of Control and Supervision of the Experiments on Animals). Protocols for experiments were approved by the Institutional Animal Ethics Committee (IAEC) of National Institute of Immunology, India, constituted by CPCSEA. Approval Code: IAEC 249/10; Approval Date: 03 August, 2010 (extended every four years).

## Data Availability Statement

The original contributions presented in this study are included in the article/supplementary material. Further inquiries can be directed to the corresponding author(s).

