## Supplementary material for "Generation of a Transgenic Mouse Model for Investigating Mitochondria in Sperm": Supplemenatry data

PGK2-MLS-GFP mice

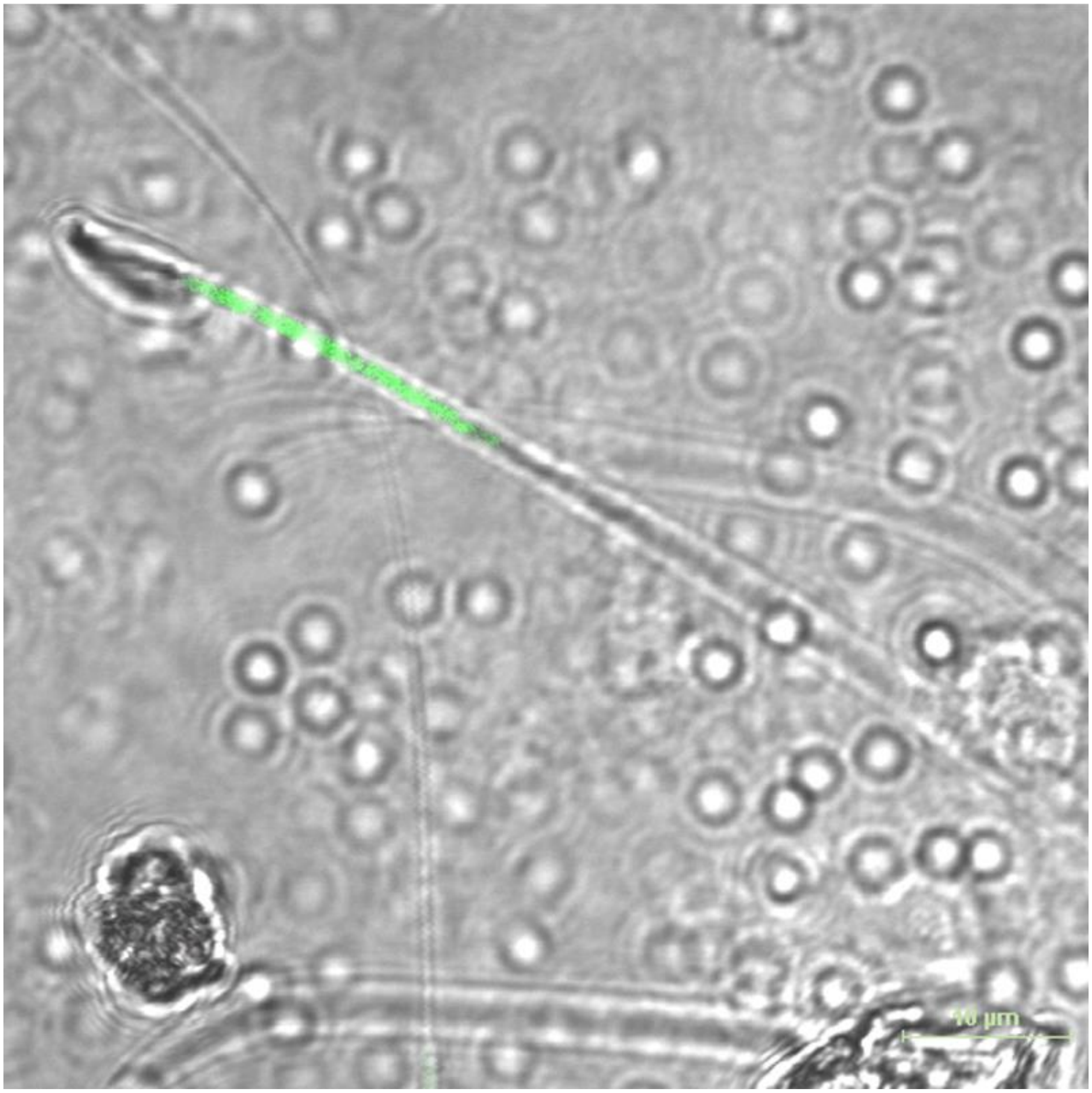


**Supplementary Figure 1**. A representative images of the spermatozoa from the PGK2-MLS-GFP transgenic mice showing the expression of GFP in the midpiece (where the mitochondria are localized).

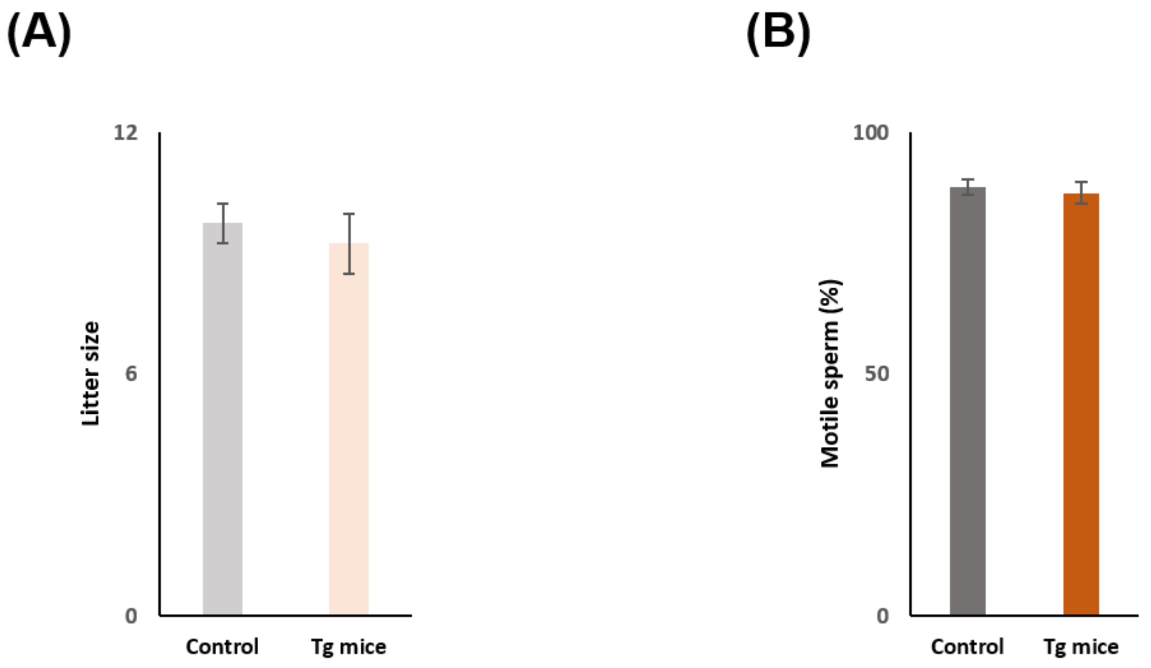


**Supplementary Figure 2**. Comparison of the litter size, sperm motility between control and PGK2-MLS-GFP transgenic mice (Tg mice). **(A)** The litter size of Wild-type (control) and PGK2-MLS-GFP transgenic mice (Tg mice). We did not observe any difference in the litter size between control and PGK2-MLS-GFP transgenic mice (Tg mice). **(B)** The motile sperm (%) of Wild-type (control) and PGK2-MLS-GFP transgenic mice (Tg mice). There was no significant difference in the motile sperm (%) between control and PGK2-MLS-GFP transgenic mice (Tg mice). $p \leq 0.05$ was considered significant.

**Supplementary Figure 3**. Full sequences of the various constructs used in this study.

CMV-MLS-EGFP

TAGTTATTAATAGTAATCAATTACGGGGTCATTAGTTCATAGCCCATATATGGAGTTCCGCGTTACATAACTTACGGTAAATGGC
CCGCCTGGCTGACCGCCCAACGACCCCCGCCCATTGACGTCAATAATGACGTATGTTCCCATAGTAACGCCAATAGGGACTTTC
CATTGACGTCAATGGGTGGAGTATTTACGGTAAACTGCCCACTTGGCAGTACATCAAGTGTATCATATGCCAAGTACGCCCCCT
ATTGACGTCAATGACGGTAAATGGCCCGCCTGGCATTATGCCCAGTACATGACCTTATGGGACTTTCCTACTTGGCAGTACATCT
ACGTATTAGTCATCGCTATTACCATGGTGATGCGGTTTTGGCAGTACATCAATGGGCGTGGATAGCGGTTTGACTCACGGGGAT
TTCCAAGTCTCCACCCCATTGACGTCAATGGGAGTTTGTTTTGGCACCAAAATCAACGGGACTTTCCAAAATGTCGTAACAACTC
CGCCCCATTGACGCAAATGGGCGGTAGGCGTGTACGGTGGGAGGTCTATATAAGCAGAGCTGGTTTAGTGAACCGTCAGATCC
GCTAGCGCTACCGGACTCAGATCTCGAGCTCAAGCTTGTGTACTCCGTGCCATCATGTCCGTCCTGACGCCGCTGCTGCTG
CGGGGCTTGACAGGCTCGGCCCGGCGGCTCCCAGTGCCGCGCGCCAAGATCCATTCGTTGCCGCCGGGGATCCACCGG
CCGGTCGCCACCATGGTGAGCAAGGGCGAGGAGCTGTTCACCGGGGTGGTGCCCATCCTGGTCGAGCTGGACGGCGACGTAA
ACGGCCACAAGTTCAGCGTGTCCGGCGAGGGCGAGGGCGATGCCACCTACGGCAAGCTGACCCTGAAGTTCATCTGCACCACC
GGCAAGCTGCCCGTGCCCTGGCCCACCCTCGTGACCACCCTGACCTACGGCGTGCAGTGCTTCAGCCGCTACCCCGACCACATG
AAGCAGCACGACTTCTTCAAGTCCGCCATGCCCGAAGGCTACGTCCAGGAGCGCACCATCTTCTTCAAGGACGACGGCAACTAC
AAGACCCGCGCCGAGGTGAAGTTCGAGGGCGACACCCTGGTGAACCGCATCGAGCTGAAGGGCATCGACTTCAAGGAGGAC
GGCAACATCCTGGGGCACAAGCTGGAGTACAACTACAACAGCCACAACGTCTATATCATGGCCGACAAGCAGAAGAACGGCAT

CAAGGTGAACTTCAAGATCCGCCACAACATCGAGGACGGCAGCGTGCAGCTCGCCGACCACTACCAGCAGAACACCCCCATCG
GCGACGGCCCCGTGCTGCTGCCCGACAACCACTACCTGAGCACCCAGTCCGCCCTGAGCAAAGACCCCAACGAGAAGCGCGAT
CACATGGTCCTGCTGGAGTTCGTGACCGCCGCCGGGATCACTCTCGGCATGGACGAGCTGTACAAGTAAAGCGGCCGCGACTC
TAGATCATAATCAGCCATACCACATTTGTAGAGGTTTTACTTGCTTTAAAAAACCTCCCACACCTCCCCCTGAACCTGAAACATAA
AATGAATGCAATTGTTGTTGTTAACTTGTTTATTGCAGCTTATAATGGTTACAAATAAAGCAATAGCATCACAAATTTCACAAAT
AAAGCATTTTTTTCACTGCATTCTAGTTGTGGTTTGTCCAAACTCATCAATGTATCTTAAGGCGTAAATTGTAAGCGTTAATATTT
TGTTAAAATTCGCGTTAAATTTTTGTTAAATCAGCTCATTTTTTAACCAATAGGCCGAAATCGGCAAAATCCCTTATAAATCAAAA
GAATAGACCGAGATAGGGTTGAGTGTTGTTCCAGTTTGGAACAAGAGTCCACTATTAAAGAACGTGGACTCCAACGTCAAAGG
GCGAAAAACCGTCTATCAGGGCGATGGCCCACTACGTGAACCATCACCCTAATCAAGTTTTTTGGGGTCGAGGTGCCGTAAAG
CACTAAATCGGAACCCTAAAGGGAGCCCCCGATTTAGAGCTTGACGGGGAAAGCCGGCGAACGTGGCGAGAAAGGAAGGGA
AGAAAGCGAAAGGAGCGGGCGCTAGGGCGCTGGCAAGTGTAGCGGTCACGCTGCGCGTAACCACCACACCCGCCGCGCTTAA
TGCGCCGCTACAGGGCGCGTCAGGTGGCACTTTTCGGGGAAATGTGCGCGGAACCCCTATTTGTTTATTTTTCTAAATACATTC
AAATATGTATCCGCTCATGAGACAATAACCCTGATAAATGCTTCAATAATATTGAAAAAGGAAGAGTCCTGAGGCGGAAAGAA
CCAGCTGTGGAATGTGTGTCAGTTAGGGTGTGGAAAGTCCCCAGGCTCCCCAGCAGGCAGAAGTATGCAAAGCATGCATCTCA
ATTAGTCAGCAACCAGGTGTGGAAAGTCCCCAGGCTCCCCAGCAGGCAGAAGTATGCAAAGCATGCATCTCAATTAGTCAGCA
ACCATAGTCCCGCCCCTAACTCCGCCCATCCCGCCCCTAACTCCGCCCAGTTCCGCCCATTCTCCGCCCCATGGCTGACTAATTTT
TTTTATTTATGCAGAGGCCGAGGCCGCCTCGGCCTCTGAGCTATTCCAGAAGTAGTGAGGAGGCTTTTTTGGAGGCCTAGGCTT
TTGCAAAGATCGATCAAGAGACAGGATGAGGATCGTTTCGCATGATTGAACAAGATGGATTGCACGCAGGTTCTCCGGCCGCT
TGGGTGGAGAGGCTATTCGGCTATGACTGGGCACAACAGACAATCGGCTGCTCTGATGCCGCCGTGTTCCGGCTGTCAGCGCA
GGGGCGCCCGGTTCTTTTTGTCAAGACCGACCTGTCCGGTGCCCTGAATGAACTGCAAGACGAGGCAGCGCGGCTATCGTGGC
TGGCCACGACGGGCGTTCCTTGCGCAGCTGTGCTCGACGTTGTCACTGAAGCGGGAAGGGACTGGCTGCTATTGGGCGAAGT
GCCGGGGCAGGATCTCCTGTCATCTCACCTTGCTCCTGCCGAGAAAGTATCCATCATGGCTGATGCAATGCGGCGGCTGCATAC
GCTTGATCCGGCTACCTGCCCATTCGACCACCAAGCGAAACATCGCATCGAGCGAGCACGTACTCGGATGGAAGCCGGTCTTG
TCGATCAGGATGATCTGGACGAAGAGCATCAGGGGCTCGCGCCAGCCGAACTGTTCGCCAGGCTCAAGGCGAGCATGCCCGA
CGGCGAGGATCTCGTCGTGACCCATGGCGATGCCTGCTTGCCGAATATCATGGTGGAAAATGGCCGCTTTTCTGGATTCATCGA
CTGTGGCCGGCTGGGTGTGGCGGACCGCTATCAGGACATAGCGTTGGCTACCCGTGATATTGCTGAAGAGCTTGGCGGCGAA
TGGGCTGACCGCTTCCTCGTGCTTTACGGTATCGCCGCTCCCGATTCGCAGCGCATCGCCTTCTATCGCCTTCTTGACGAGTTCTT
CTGAGCGGGACTCTGGGGTTCGAAATGACCGACCAAGCGACGCCCAACCTGCCATCACGAGATTTCGATTCCACCGCCGCCTTC
TATGAAAGGTTGGGCTTCGGAATCGTTTTCCGGGACGCCGGCTGGATGATCCTCCAGCGCGGGGATCTCATGCTGGAGTTCTT
CGCCCACCCTAGGGGGAGGCTAACTGAAACACGGAAGGAGACAATACCGGAAGGAACCCGCGCTATGACGGCAATAAAAAG
ACAGAATAAAACGCACGGTGTTGGGTCGTTTGTTCATAAACGCGGGGTTCGGTCCCAGGGCTGGCACTCTGTCGATACCCCAC
CGAGACCCCATTGGGGCCAATACGCCCGCGTTTCTTCCTTTTCCCCACCCCACCCCCCAAGTTCGGGTGAAGGCCCAGGGCTCG
CAGCCAACGTCGGGGCGGCAGGCCCTGCCATAGCCTCAGGTTACTCATATATACTTTAGATTGATTTAAAACTTCATTTTTAATT
TAAAAGGATCTAGGTGAAGATCCTTTTTGATAATCTCATGACCAAAATCCCTTAACGTGAGTTTTCGTTCCACTGAGCGTCAGAC
CCCGTAGAAAAGATCAAAGGATCTTCTTGAGATCCTTTTTTTCTGCGCGTAATCTGCTGCTTGCAAACAAAAAAACCACCGCTAC
CAGCGGTGGTTTGTTTGCCGGATCAAGAGCTACCAACTCTTTTTCCGAAGGTAACTGGCTTCAGCAGAGCGCAGATACCAAATA
CTGTCCTTCTAGTGTAGCCGTAGTTAGGCCACCACTTCAAGAACTCTGTAGCACCGCCTACATACCTCGCTCTGCTAATCCTGTTA
CCAGTGGCTGCTGCCAGTGGCGATAAGTCGTGTCTTACCGGGTTGGACTCAAGACGATAGTTACCGGATAAGGCGCAGCGGTC
GGGCTGAACGGGGGGTTCGTGCACACAGCCCAGCTTGGAGCGAACGACCTACACCGAACTGAGATACCTACAGCGTGAGCTA
TGAGAAAGCGCCACGCTTCCCGAAGGGAGAAAGGCGGACAGGTATCCGGTAAGCGGCAGGGTCGGAACAGGAGAGCGCAC
GAGGGAGCTTCCAGGGGGAAACGCCTGGTATCTTTATAGTCCTGTCGGGTTTCGCCACCTCTGACTTGAGCGTCGATTTTTGTG
ATGCTCGTCAGGGGGGCGGAGCCTATGGAAAAACGCCAGCAACGCGGCCTTTTTACGGTTCCTGGCCTTTT
GCTGGCCTTTTGCTCACATGTTCTTTCCTGCGTTATCCCCTGATTCTGTGGATAACCGTATTACCGC
CATGCAT

### PGK2-MLS-EGFPN2

TAGTTATTAATCTAGAAATAGGATTGATGACTTGTATGGTAAGTCTATGCATATTTCT
AAGAAATTGCCAAGCTGTTTCCCAAAATGTCTGTAGCATATTTCATTCCTGGGAGCA

ACAAATGAGAGTTTCTGTTACTATATATTTTAACACCATTTAAGTTTGCTTTTAAATG
CTGTTGACTCGATGTTCAACATTGAGGAACCGATAAACTGTTTTTTAAAGTGGCTGC
ACCATTAGACATTCCTATCAGCAATAAATGAGGATTCCAATTTGTCCACTTCTTTCTC
AACACTTGTTTTTATTTTTTATTTTAGCCATCTTAATAGGAGATGCCTCAAATATATTT
CTGTCAAAGCCTTCATTTTACATATTTAGTTCACTCAACTTATGGAGTAGAATGAAAA
AAACTGTAAAGCCAATTTTCATTACTAAACCAACTAGCCACATACATTTGCTTTCTCC
AAAAATAGAATATCTTACTCACTTTCAATTCTAATATATGTTTTCATCTTCATGAAAA
CCATGCATTTATTTTTCAGCTCAATTTCTAAGTTGTACAGGTAGGAAAGCAAACAGA
GAGGAGACAAAAAATAAGGAGCCTCATCATAACATTGTAGTCTGAAAGAAATGTCA
CCATTACTTTCAGCACTTGTTACTGATACTCTCTTTGCTATGATCTTATGGATTCTCC
TGCAGAAGTGACCAAACATTCTTTCATGGTAGTCTGAAAATGGAAGAAGCAAAGTA
ATGAAAATATGACAGCATTCCCACCACGTCCCAGAATAGATCTAAAGATGAAGACT
AACAACAAGCCACATCCTCAAAGAAATATTTATCAAGACAAGGAAAAAACAGAACT
ATACTATCCTGACTTGAATAAGCAAGGTTTCATATGAAAATCATTATTTCAAATATTC
CTTTTATTTGGCACTGCAGAGGTTTTTACATATCAAAATGGTTAAGATTTGACATGAA
TGAGGTGTATGTAGGTTTGCGGGGGTGGGGGTGGAGTTCTTTTTGTTTTGTTTTTAAT
AATAAAAGCAACTGTTAACCGAGCTGTGGGGTGGGGGCAAAAAGAGCCAGAAGGCG
GCGCACACCTCAGGACTATTCTTGTTTTTTTAGAACCATTCTTATTCTGGGGTTTCTTC
ACCTACCCAAGTCTCGCCTGAAGCCAGGTACAGCTCTATTCCACTACATGACCCTCT
GCCCAGGAAGTTGGAATCTTCACCTAGCAACACAGTTCAGATCGAGATTGACAGGAC
CATGAGCCAATCACAAAGCTAGATTTGCTTTCCAGTCTAACAGTGGCCGTTGTGCTG
GAGACAGTGAGGAGAAGAAAGGGGCGGGACAAGGGCAAAGGCGTTAGAAGTCACC
ACCGACCCAGCCCCTCAACAGCAAGTTGGTTCTTCAGCATTAAGATCCAGGTGTCAG
CCTATGTCTTTATATTGTCAAGCTCGAGCTCAAGCTTGTGTACTCCGTGCCATCATGTCCG
TCCTGACGCCGCTGCTGCTGCGGGGCTTGACAGGCTCGGCCCGGCGGCTCCCAGTGCCGCGCGCC
AAGATCCATTCGTTGCCGCCGGGGATCCACCGGCCGGTCGCCACCATGGTGAGCAAGGG
CGAGGAGCTGTTCACCGGGGTGGTGCCCATCCTGGTCGAGCTGGACGGCGACGTAA
ACGGCCACAAGTTCAGCGTGTCCGGCGAGGGCGAGGGCGATGCCACCTACGGCAAG
CTGACCCTGAAGTTCATCTGCACCACCGGCAAGCTGCCCGTGCCCTGGCCCACCCTC
GTGACCACCCTGACCTACGGCGTGCAGTGCTTCAGCCGCTACCCCGACCACATGAAG
CAGCACGACTTCTTCAAGTCCGCCATGCCCGAAGGCTACGTCCAGGAGCGCACCATC
TTCTTCAAGGACGACGGCAACTACAAGACCCGCGCCGAGGTGAAGTTCGAGGGCGA
CACCCTGGTGAACCGCATCGAGCTGAAGGGCATCGACTTCAAGGAGGACGGCAACA
TCCTGGGGCACAAGCTGGAGTACAACTACAACAGCCACAACGTCTATATCATGGCCG
ACAAGCAGAAGAACGGCATCAAGGTGAACTTCAAGATCCGCCACAACATCGAGGAC
GGCAGCGTGCAGCTCGCCGACCACTACCAGCAGAACACCCCCATCGGCGACGGCCC
CGTGCTGCTGCCCGACAACCACTACCTGAGCACCCAGTCCGCCCTGAGCAAAGACCC
CAACGAGAAGCGCGATCACATGGTCCTGCTGGAGTTCGTGACCGCCGCCGGGATCAC
TCTCGGCATGGACGAGCTGTACAAGTAAAGCGGCCGCGACTCTAGATCATAATCAGC
CATACCACATTTGTAGAGGTTTTACTTGCTTTAAAAAACCTCCCACACCTCCCCCTGA
ACCTGAAACATAAAATGAATGCAATTGTTGTTGTTAACTTGTTTATTGCAGCTTATAA
TGGTTACAAATAAAGCAATAGCATCACAAATTTCACAAATAAAGCATTTTTTTCACT
GCATTCTAGTTGTGGTTTGTCCAAACTCATCAATGTATCTTAAGGCGTAAATTGTAAG
CGTTAATATTTTGTTAAAATTCGCGTTAAATTTTTGTTAAATCAGCTCATTTTTTAACC
AATAGGCCGAAATCGGCAAAATCCCTTATAAATCAAAAGAATAGACCGAGATAGGG
TTGAGTGTTGTTCCAGTTTGGAACAAGAGTCCACTATTAAAGAACGTGGACTCCAAC

GTCAAAGGGCGAAAAACCGTCTATCAGGGCGATGGCCCACTACGTGAACCATCACC
CTAATCAAGTTTTTTGGGGTCGAGGTGCCGTAAAGCACTAAATCGGAACCCTAAAGG
GAGCCCCCGATTTAGAGCTTGACGGGGAAAGCCGGCGAACGTGGCGAGAAAGGAAG
GGAAGAAAGCGAAAGGAGCGGGCGCTAGGGCGCTGGCAAGTGTAGCGGTCACGCTG
CGCGTAACCACCACACCCGCCGCGCTTAATGCGCCGCTACAGGGCGCGTCAGGTGGC
ACTTTTCGGGGAAATGTGCGCGGAACCCCTATTTGTTTATTTTTCTAAATACATTCAA
ATATGTATCCGCTCATGAGACAATAACCCTGATAAATGCTTCAATAATATTGAAAAA
GGAAGAGTCCTGAGGCGGAAAGAACCAGCTGTGGAATGTGTGTCAGTTAGGGTGTG
GAAAGTCCCCAGGCTCCCCAGCAGGCAGAAGTATGCAAAGCATGCATCTCAATTAGT
CAGCAACCAGGTGTGGAAAGTCCCCAGGCTCCCCAGCAGGCAGAAGTATGCAAAGC
ATGCATCTCAATTAGTCAGCAACCATAGTCCCGCCCCTAACTCCGCCCATCCCGCCCC
TAACTCCGCCCAGTTCCGCCCATTCTCCGCCCCATGGCTGACTAATTTTTTTTATTTAT
GCAGAGGCCGAGGCCGCCTCGGCCTCTGAGCTATTCCAGAAGTAGTGAGGAGGCTTT
TTTGGAGGCCTAGGCTTTTGCAAAGATCGATCAAGAGACAGGATGAGGATCGTTTCG
CATGATTGAACAAGATGGATTGCACGCAGGTTCTCCGGCCGCTTGGGTGGAGAGGCT
ATTCGGCTATGACTGGGCACAACAGACAATCGGCTGCTCTGATGCCGCCGTGTTCCG
GCTGTCAGCGCAGGGGCGCCCGGTTCTTTTTGTCAAGACCGACCTGTCCGGTGCCCT
GAATGAACTGCAAGACGAGGCAGCGCGGCTATCGTGGCTGGCCACGACGGGCGTTC
CTTGCGCAGCTGTGCTCGACGTTGTCACTGAAGCGGGAAGGGACTGGCTGCTATTGG
GCGAAGTGCCGGGGCAGGATCTCCTGTCATCTCACCTTGCTCCTGCCGAGAAAGTAT
CCATCATGGCTGATGCAATGCGGCGGCTGCATACGCTTGATCCGGCTACCTGCCCAT
TCGACCACCAAGCGAAACATCGCATCGAGCGAGCACGTACTCGGATGGAAGCCGGT
CTTGTCGATCAGGATGATCTGGACGAAGAGCATCAGGGGCTCGCGCCAGCCGAACT
GTTCGCCAGGCTCAAGGCGAGCATGCCCGACGGCGAGGATCTCGTCGTGACCCATGG
CGATGCCTGCTTGCCGAATATCATGGTGGAAAATGGCCGCTTTTCTGGATTCATCGA
CTGTGGCCGGCTGGGTGTGGCGGACCGCTATCAGGACATAGCGTTGGCTACCCGTGA
TATTGCTGAAGAGCTTGGCGGCGAATGGGCTGACCGCTTCCTCGTGCTTTACGGTAT
CGCCGCTCCCGATTCGCAGCGCATCGCCTTCTATCGCCTTCTTGACGAGTTCTTCTGA
GCGGGACTCTGGGGTTCGAAATGACCGACCAAGCGACGCCCAACCTGCCATCACGA
GATTTCGATTCCACCGCCGCCTTCTATGAAAGGTTGGGCTTCGGAATCGTTTTCCGGG
ACGCCGGCTGGATGATCCTCCAGCGCGGGGATCTCATGCTGGAGTTCTTCGCCCACC
CTAGGGGGAGGCTAACTGAAACACGGAAGGAGACAATACCGGAAGGAACCCGCGCT
ATGACGGCAATAAAAAGACAGAATAAAACGCACGGTGTTGGGTCGTTTGTTCATAA
ACGCGGGGTTCGGTCCCAGGGCTGGCACTCTGTCGATACCCCACCGAGACCCCATTG
GGGCCAATACGCCCGCGTTTCTTCCTTTTCCCCACCCCACCCCCCAAGTTCGGGTGAA
GGCCCAGGGCTCGCAGCCAACGTCGGGGCGGCAGGCCCTGCCATAGCCTCAGGTTA
CTCATATATACTTTAGATTGATTTAAAACTTCATTTTTAATTTAAAAGGATCTAGGTG
AAGATCCTTTTTGATAATCTCATGACCAAAATCCCTTAACGTGAGTTTTCGTTCCACT
GAGCGTCAGACCCCGTAGAAAAGATCAAAGGATCTTCTTGAGATCCTTTTTTTCTGC
GCGTAATCTGCTGCTTGCAAACAAAAAAACCACCGCTACCAGCGGTGGTTTGTTTGC
CGGATCAAGAGCTACCAACTCTTTTTCCGAAGGTAACTGGCTTCAGCAGAGCGCAGA
TACCAAATACTGTCCTTCTAGTGTAGCCGTAGTTAGGCCACCACTTCAAGAACTCTGT
AGCACCGCCTACATACCTCGCTCTGCTAATCCTGTTACCAGTGGCTGCTGCCAGTGGC
GATAAGTCGTGTCTTACCGGGTTGGACTCAAGACGATAGTTACCGGATAAGGCGCAG
CGGTCGGGCTGAACGGGGGGTTCGTGCACACAGCCCAGCTTGGAGCGAACGACCTA
CACCGAACTGAGATACCTACAGCGTGAGCTATGAGAAAGCGCCACGCTTCCCGAAG
GGAGAAAGGCGGACAGGTATCCGGTAAGCGGCAGGGTCGGAACAGGAGAGCGCAC

GAGGGAGCTTCCAGGGGGAAACGCCTGGTATCTTTATAGTCCTGTCGGGTTTCGCCA
CCTCTGACTTGAGCGTCGATTTTTGTGATGCTCGTCAGGGGGGCGGAGCCTATGGAA
AAACGCCAGCAACGCGGCCTTTTTACGGTTCCTGGCCTTTTGCTGGCCTTTTGCTCAC
ATGTTCTTTCCTGCGTTATCCCCTGATTCTGTGGATAACCGTATTACCGCCATGCAT
